# TensorPLS: an R package for exploring molecular and temporal patterns in longitudinal omics data using PLS-DA

**DOI:** 10.64898/2026.11.03.763745

**Authors:** Alessandro Giordano, Ana Conesa, Leandro Balzano-Nogueira

## Abstract

Longitudinal omics datasets combine high-dimensional repeated measurements with missing observations, complicating supervised modelling and temporal interpretation. We present TensorPLS, an R package integrating missing-data handling, matrix-based partial least squares discriminant analysis (PLS-DA), structured variable importance, subject-level nested cross-validation and downstream functional enrichment. Data are represented as subjects × features × time tensors. Missing values are handled by Tucker3 reconstruction or feature–time mean imputation, after which tensors are unfolded along the subject mode for PLS-DA. Cumulative variable importance in projection (VIP) scores are mapped to feature–time coordinates and summarised across time and cumulative PLS models. During nested cross-validation, imputation, component tuning, VIP-based feature selection and refitting are performed within training partitions. Full-cohort models are fitted separately for descriptive interpretation. We evaluated TensorPLS on two public longitudinal transcriptomic datasets using five-by-five nested cross-validation. In GSE68310 (109 subjects), Tucker3 reconstruction yielded 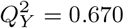, AUROC 0.971 and balanced accuracy 0.931. In GSE182616 (66 subjects), feature–time mean imputation yielded 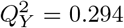, AUROC 0.827 and balanced accuracy 0.756. Predictive metrics were calculated from pooled outer held-out predictions. TensorPLS supports predictive assessment, functional enrichment of selected genes and temporal interpretation of model-assigned feature importance.

## Introduction

Longitudinal omics studies measure molecular profiles from the same individuals at multiple time points, allowing molecular changes to be examined in relation to disease progression, treatment response, and other evolving biological processes (Taheriyoun et al., 2026). These datasets are typically high-dimensional and may also contain missing measurements across time, complicating both modelling and interpretation (Taheriyoun et al., 2026). When the same molecular features are measured over a common set of time points, the data can naturally be represented as a three-way array with dimensions subjects × features × time (Kolda and Bader, 2009; Kodikara et al., 2026). This representation preserves the temporal organisation of the measurements and, in particular, the correspondence between each molecular feature and the time points at which it was observed.

Several multivariate approaches have been developed for multiway data. N-way partial least squares (N-PLS), for example, extends PLS regression to predictor arrays with three or more modes (Bro, 1996), while tensor decompositions such as Tucker3 models provide a general representation of multiway variation (Kolda and Bader, 2009). These methods have also been applied to the integration of multiway omics data (Conesa et al., 2010). More recently, tensorOmics introduced tensor-based extensions of PCA and PLS for longitudinal single- and multi-omics data, including supervised tensor PLS discriminant analysis and multi-block formulations (Kodikara et al., 2026). These approaches retain the multiway structure during model fitting. Direct tensor modelling, however, is not the only way to make use of longitudinal structure. Different stages of an analysis can operate on different representations: tensor methods can be used where the multiway organisation is directly relevant, whereas established matrix-based supervised methods can be applied after unfolding the data along the subject mode. In this representation, each feature–time combination becomes a predictor while its mapping to the original tensor dimensions can be retained for interpretation (Kolda and Bader, 2009). A practical challenge is therefore not only how to model longitudinal tensors, but how to combine missing-data reconstruction, supervised modelling, feature selection, and validation without losing this structure or allowing information from held-out observations to influence model development (Varma and Simon, 2006). The general analytical concept underlying TensorPLS was inspired by our previous work on longitudinal omics data in the TEDDY study (Balzano-Nogueira et al., 2021).

Here, we present TensorPLS, an R package that integrates missing-data handling, supervised modelling, feature selection and predictive validation into a single workflow for longitudinal omics data. The package also provides functional enrichment of selected genes, supporting downstream biological interpretation.

## Methods

### TensorPLS overview

TensorPLS provides a unified workflow for supervised analysis of longitudinal omics data while preserving the correspondence between molecular features and measurement times (Fig. 1). Missing values can be handled either through Tucker3 reconstruction of the three-way array or through feature–time mean imputation. TensorPLS implements a matrix-based PLS model using the NIPALS algorithm. The completed tensor is unfolded along the subject mode, while the original feature–time coordinates are retained for subsequent VIP-based interpretation.

**Figure 1.**
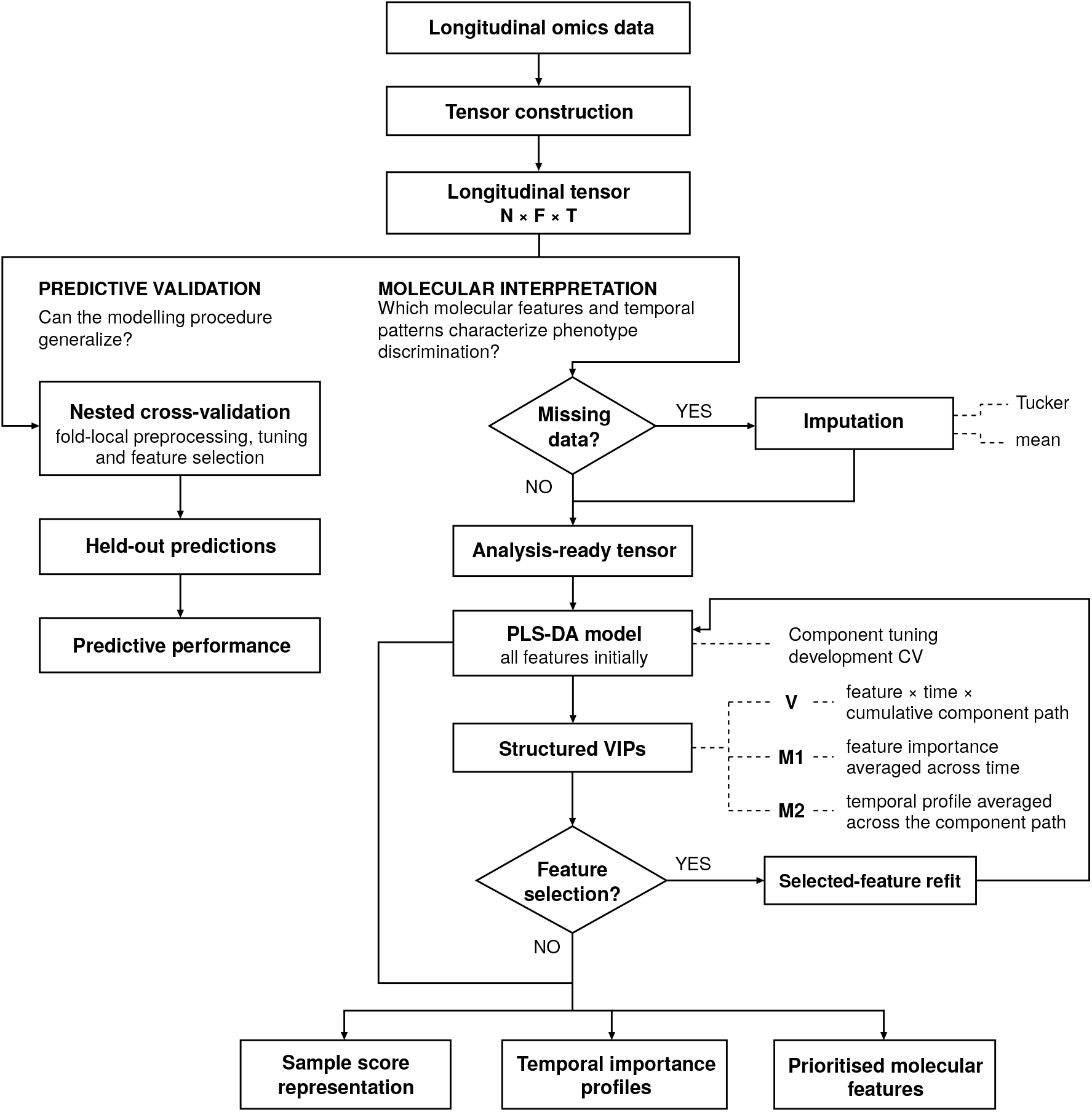
Overview of the TensorPLS workflow. Longitudinal omics data are organised as a tensor with dimensions subjects *×* features *×* time. Predictive performance is evaluated through nested cross-validation with training-local preprocessing and model development, while the full-cohort analysis provides sample score representations, temporal importance profiles and prioritised molecular features for interpretation. Missing data can be handled using Tucker3 reconstruction or feature–time mean imputation. **Alt text:** Flowchart of TensorPLS. A longitudinal data tensor branches into nested cross-validation for held-out performance and full-cohort analysis for interpretation, including optional imputation, PLS-DA, VIP selection, and molecular and temporal summaries.

The mapping between unfolded predictors and their original feature–time coordinates is retained throughout the analysis, allowing variable-importance estimates from the PLS model to be reorganised into longitudinal representations. Full-cohort modelling provides sample score representations, temporal importance profiles and prioritised molecular features to characterise patterns associated with class discrimination. Separately, nested cross-validation evaluates the out-of-sample predictive performance of the complete modelling pipeline, with preprocessing, component selection, feature selection and refitting performed within the corresponding training partitions.

### Study datasets and preprocessing

TensorPLS was evaluated on two publicly available longitudinal transcriptomic datasets, GSE68310 and GSE182616. For both datasets, processed expression measurements deposited in the Gene Expression Omnibus (GEO) were used without additional normalisation or expression-based feature filtering. Samples were organised into subjects × features × time tensors, retaining the correspondence between each subject, sampling time and microarray probe.

#### GSE68310: respiratory viral infections

GSE68310 (Zhai et al., 2015) contains longitudinal blood gene-expression measurements generated using the Illumina HumanHT-12 v4 microarray platform (GPL10558). Expression values were obtained from the deposited GEO Series Matrix, while respiratory viral diagnoses were derived from the accompanying subject-level PCR metadata.

Subjects were classified as influenza-positive when influenza was detected by PCR, including coinfections, and as other virus-positive when at least one non-influenza respiratory virus was detected in the absence of influenza. Subjects without a detected virus were excluded, leaving 109 subjects: 73 influenza-positive and 36 other virus-positive.

Measurements were organised at four sampling times (Day0, Day2, Day4 and Day6). After excluding technical control probes, the resulting tensor had dimensions 109 × 47,162 × 4. Of the 436 expected subject–time profiles, 21 were absent (4.82%), corresponding to 990,402 missing tensor entries. Tucker3-based reconstruction was used to handle missing data.

#### GSE182616: burn-injury severity

GSE182616 (Keyloun et al., 2022) contains longitudinal whole-blood gene-expression measurements obtained using the Agilent microarray platform GPL17077. The deposited values represent expression log ratios between patient RNA and an Agilent Universal Human Reference. These processed measurements were used as deposited.

The analysis was restricted to samples collected at 0, 2, 4, 8, 12 and 24 hours. Two subjects without measurements at these time points were excluded, leaving 66 subjects. Burn severity was defined according to total body surface area (TBSA), with 34 subjects classified as Severe (TBSA ≥ 20%) and 32 as LessSevere (TBSA *<* 20%).

After excluding technical control probes, the resulting tensor had dimensions 66 × 50,599 × 6. Of the 396 expected subject– time profiles, 91 were absent (22.98%). Feature–time mean imputation was used to handle missing data.

### Missing-data handling

In both applications, missingness occurred at the visit level, with entire subject–time expression profiles unavailable. Tucker3-based reconstruction was used for GSE68310, whereas feature–time mean imputation was used for GSE182616.

Imputation was performed separately for predictive validation and full-cohort interpretation. During nested cross-validation, imputation parameters were estimated from training subjects and subsequently applied to held-out subjects. For full-cohort interpretation, missing values were imputed using all available subjects before PLS-DA modelling, VIP-based feature selection and visualisation. Full-cohort imputation was not used to estimate predictive performance.

#### Tucker3-based reconstruction of missing data

Tucker3 reconstruction preserves the three-way organisation of the longitudinal data during missing-data estimation. Let

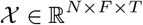

denote a subjects × features × time tensor, and let Ω denote the set of its observed entries.

Feature–time means are estimated from the available observations and used to centre the tensor,

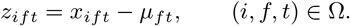

The centred tensor is represented through a Tucker3 decomposition,

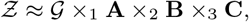

where G is the core tensor and **A, B** and **C** contain the factors associated with the subject, feature and time modes, respectively.

The Tucker3 representation is estimated from the observed entries by minimising

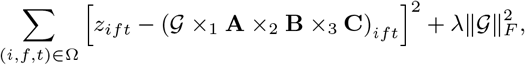

where *λ* controls a ridge penalty on the Tucker3 core. Missing entries do not contribute to the observed-entry objective. After fitting, originally observed measurements are retained unchanged and only missing entries are replaced by their fitted low-rank reconstructions.

##### Predictive train/apply reconstruction

During predictive validation, Tucker3 reconstruction is instead estimated within each training partition. The Tucker3 core, feature- and time-mode factors and training-derived centering quantities are learned from the training tensor and kept fixed when the fitted representation is applied to held-out subjects. For each held-out subject, subject-specific latent coordinates are estimated from its available measurements using observed-entry least squares and are then used to reconstruct its missing entries. Held-out measurements therefore contribute to estimation of that subject’s coordinates but do not re-estimate the shared Tucker3 representation learned from the training partition.

For GSE68310, Tucker3 ranks were fixed to (4, 4, 3), with a ridge coefficient of *λ* = 0.001, a convergence tolerance of 10^−6^ and a maximum of 2,000 iterations. HOSVD initialization and a projection tolerance of 10^−10^ were used, with random seed 123.

#### Feature–time mean imputation

As a simpler alternative, missing values can be replaced by the mean of the observed measurements at the corresponding feature–time position,

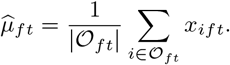

Observed values are retained unchanged. For full-cohort modelling, the means are estimated using all available subjects. During predictive validation, they are estimated independently within each training partition and the training-derived values are subsequently applied to the corresponding held-out subjects.

### PLS-DA modelling

After missing-data handling, the completed subject-mode unfolding is analysed using a matrix-based NIPALS PLS model (Wold et al., 2001), with class-encoded responses used for discriminant analysis (Barker and Rayens, 2003).

For binary classification, the outcome is represented as a single 0*/*1 response. Predictor columns and the encoded response are centred and scaled using column means and sample standard deviations estimated from the training data. Constant predictor columns are retained, with their scaling factor set to one.

For component *a*, the predictor score is obtained as

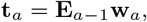

where **E**_*a*−1_ is the predictor residual matrix after extraction of the preceding components and **w**_*a*_ is the normalised predictor weight vector. Predictor and response residual matrices are successively deflated, yielding a sequence of latent components associated with the encoded response.

A model containing *H* components corresponds to the cumulative PLS path formed by components 1, …, *H*. During nested predictive evaluation, *H* is treated as a tuning parameter and is selected using inner resampling restricted to the corresponding outer-training subjects.

Predictions for held-out subjects are generated using the coefficients and scaling parameters estimated from the training data. For a single binary response, continuous PLS predictions are converted to class assignments using the fixed midpoint threshold of 0.5, with values at or above the threshold assigned to the positive class. The threshold is not tuned during model development. Continuous predictions are used as discrimination scores and are not interpreted as calibrated probabilities.

### Structured variable importance

Variable importance in projection (VIP) has been extended to N-PLS models (Favilla et al., 2013) and used for longitudinal omics analysis in the earlier TEDDY study (Balzano-Nogueira et al., 2021). In TensorPLS, variable importance is calculated for each feature–time predictor in the unfolded PLS model. Let

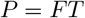

denote the total number of unfolded predictors, **w**_*a*_ the normalised predictor weights for component *a*, and **t**_*a*_ the corresponding PLS score.

The response contribution associated with component *a* is defined as

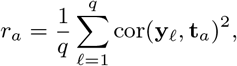

where *q* denotes the number of encoded response columns. For the binary analysis considered here, *q* = 1.

For unfolded predictor *j*, the cumulative VIP associated with the model containing the first *h* PLS components is

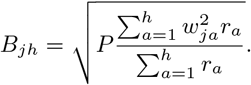

The index *h* therefore denotes the cumulative model containing components 1, …, *h*, rather than the isolated contribution of component *h*.

Using the feature–time mapping defined during unfolding, the cumulative VIP matrix is reshaped as

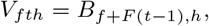

forming the structured representation

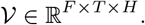

The three dimensions of V retain complementary information from the fitted PLS model. For a given molecular feature, *V*_*fth*_ describes its importance at measurement time *t* for the cumulative model containing components 1, …, *h*. Thus, V preserves both the longitudinal location of a predictor and the evolution of its importance as additional PLS components are included.

Two complementary marginal summaries are derived from V. The first marginal summary averages over measurement times,

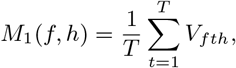

yielding an *F* × *H* representation. It summarizes the overall importance of each molecular feature across the longitudinal window as model complexity increases along the cumulative PLS path.

The second marginal summary averages over the cumulative component path,

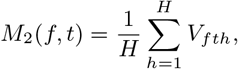

yielding an *F* × *T* representation. It preserves the temporal dimension and therefore describes how the model-assigned importance of each molecular feature is distributed across measurement times.

The full representation *V* and the marginal summaries *M*_1_ and *M*_2_ are derived from the same underlying cumulative VIP estimates and are therefore interpreted as complementary views of variable importance rather than independent VIP algorithms.

### VIP-based feature selection

VIP-guided feature selection was also used in the earlier TEDDY analysis (Balzano-Nogueira et al., 2021). In TensorPLS, feature selection is based on the structured VIP representations and is performed within training partitions during predictive validation or on the complete cohort for molecular interpretation. For a given candidate component count *H*, an initial PLS model is fitted using all molecular features, and its VIP values are used to define the retained feature set before refitting the model on the reduced predictor matrix.

TensorPLS supports percentile-based selection from individual VIP representations or from their intersections. In the unfolded cumulative VIP representation, thresholds are calculated across the *FT* feature–time predictors for each cumulative component column. A molecular feature is retained if at least one of its feature–time predictors reaches or exceeds the corresponding percentile criterion.

When multiple VIP representations are combined by intersection, the final feature set contains only molecular features retained by each contributing representation. All time-point predictors of a retained molecular feature are then included in the reduced model, which is refitted using the same *H*. During predictive validation, VIP calculation, threshold estimation, feature selection and refitting are repeated independently within each training partition. For both applications reported here, feature selection used the intersection between percentile-based selections from the unfolded cumulative VIP representation and the time-resolved *M*_2_ representation, with a fixed 99th-percentile threshold. Neither the selection rule nor the percentile threshold was tuned during predictive validation.

We additionally compared the four VIP-based selection strategies (*V, M*_1_, *M*_2_ and *V* ∩ *M*_2_) using the same nested cross-validation partitions. Predictive performance and feature-selection agreement across outer folds were evaluated as described in Supplementary Methods, Section S6.

### Predictive validation and full-cohort interpretation

TensorPLS supports two complementary analytical objectives. Full-cohort modelling is used to characterise class-associated molecular and temporal patterns, whereas nested cross-validation evaluates the out-of-sample performance of the complete predictive pipeline.

Predictive performance was evaluated using subject-stratified nested cross-validation with five outer folds and five inner folds. Within each outer-training partition, missing-data handling, predictor centering and scaling, PLS component selection, VIP-based feature selection and model refitting were performed using training data only. Candidate component counts *H* = 1, …, 10 were evaluated through inner resampling, with component selection based on the validation performance of the complete feature-selection and refitting pipeline. The selected pipeline was then refitted on the complete outer-training set and applied to the corresponding outer-test subjects. Pooled outer predictions were used to calculate AUROC, balanced accuracy, sensitivity, specificity, BER and 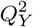. For binary classification, the fixed midpoint threshold of 0.5 was used to convert continuous PLS predictions into class assignments.

For full-cohort molecular interpretation, the component count was selected through five-fold cross-validation of the VIP-based selection and refitting pipeline, using the fixed *V* ∩ *M*_2_ strategy and 99th-percentile threshold. An all-feature PLS model was then fitted on the imputed full cohort, followed by VIP-based feature selection and refitting with the selected component count. Sample scores, temporal importance profiles and prioritised molecular features were obtained for descriptive interpretation. Out-of-sample predictive performance was evaluated separately through nested cross-validation.

### Functional enrichment and temporal interpretation

TensorPLS provides a gene-based functional enrichment module that accepts selected genes and a user-defined background as Entrez identifiers or gene symbols. Gene symbols are converted to Entrez identifiers through unambiguous mappings using org.Hs.eg.db.

The module performs over-representation analysis using GO Biological Process, Reactome and MSigDB Hallmark (Liberzon et al., 2015) gene sets, through clusterProfiler (Yu et al., 2012) and ReactomePA. Enrichment is evaluated using the hypergeometric test, with Benjamini–Hochberg correction applied separately to each database. Gene-set sizes were restricted to 10–500 genes within the effective annotation background, and adjusted *P <* 0.05 was used to define significance.

For the two microarray applications, selected probes were mapped to Entrez identifiers using platform-specific annotations. The background comprised unique, recognised genes represented among all analysed probes. Hallmark analysis used MSigDB version 2026.1.Hs.

For temporal interpretation, selected probes associated with enriched gene sets were linked to their *M*_2_ profiles from the full-cohort all-feature model. Profiles were summarised using the median and interquartile range across probes.

## Results

### Applications to longitudinal transcriptomic datasets GSE68310: respiratory viral infections

We first applied TensorPLS to GSE68310, comprising 109 subjects with PCR-confirmed respiratory viral infections. The outcome distinguished 73 influenza-positive subjects, including coinfections, from 36 subjects positive for other respiratory viruses but negative for influenza. The input tensor contained 47,162 probes measured at Day0, Day2, Day4 and Day6. Of the 436 expected subject–time profiles, 21 were entirely missing (4.82%), corresponding to 990,402 missing tensor entries.

Missing values were handled using Tucker3 reconstruction with fixed ranks (4, 4, 3), fitted separately within training partitions during predictive validation. Five-by-five subject-level nested cross-validation yielded 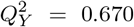, an AUROC of 0.971 and a balanced accuracy of 0.931 from pooled outer held-out predictions. Sensitivity and specificity were 0.945 and 0.917, respectively. Across the five outer folds, the selected number of PLS components ranged from 1 to 8 and the number of retained probes ranged from 1,073 to 1,149.

For full-cohort interpretation, component development selected *H* = 5. VIP-based selection retained 1,132 probes, and the reduced model was refitted with the same component count. The selected-feature score representation is shown in Fig. 2A. The mean temporal *M*_2_ profile shows the average importance assigned to selected probes at each sampling time. Mean importance was highest at Day4, closely followed by Day2, and lower at Day0 and Day6 (Fig. 2B).

**Figure 2.**
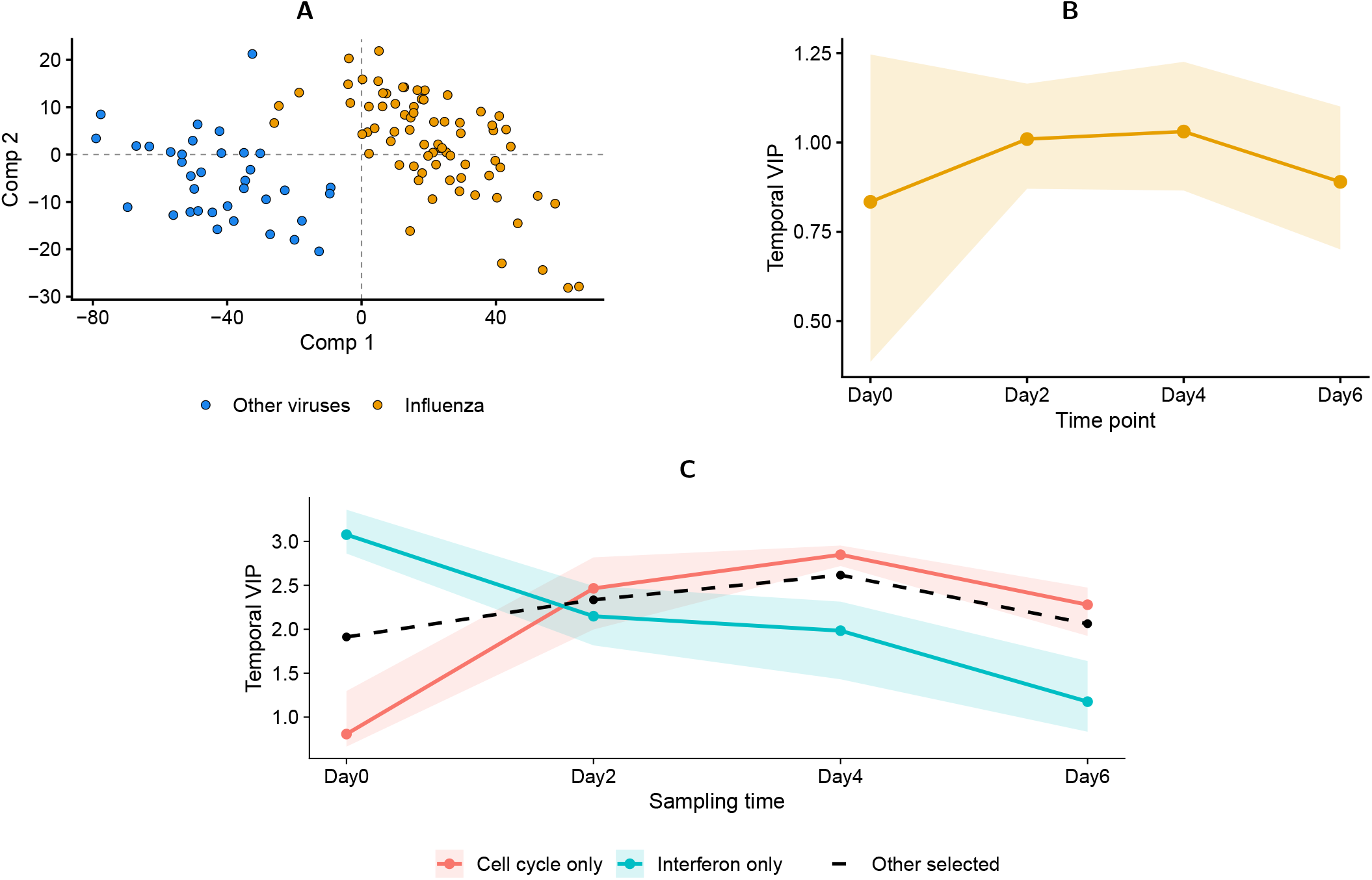
Full-cohort descriptive analysis of GSE68310. (A) Sample scores from the model refitted on 1,132 selected probes. (B) Mean temporal *M*_2_ profile from the selected-feature model. (C) Median *M*_2_ profiles of interferon-associated (83 probes) and cell-cycle-associated (81 probes) features from the all-feature model. Shared genes were excluded to define disjoint groups. The dashed line represents the remaining 966 selected probes. Shaded areas in (B) and (C) indicate interquartile ranges across probes, not confidence intervals. None of the panels shows held-out predictive results. **Alt text:** Panel A plots full-cohort PLS-DA scores by infection class. Panel B plots average temporal VIP, highest at Day4. Panel C compares interferon-associated and cell-cycle-associated VIP profiles, peaking at Day0 and Day4, respectively.

Functional enrichment identified 306 significant GO Biological Process terms, 55 Reactome pathways and seven Hallmark gene sets, including interferon responses, E2F targets and G2M checkpoint.

After excluding shared genes, median *M*_2_ importance peaked at Day0 for 83 interferon-associated probes (3.08) and at Day4 for 81 cell-cycle-associated probes (2.85; Fig. 2C). These profiles illustrate how functionally distinct subsets within the same selected signature can exhibit different patterns of model-assigned importance across sampling times, providing information that is not captured by enrichment results or an aggregate temporal VIP profile alone.

#### GSE182616: burn-injury severity

We next analysed GSE182616, comprising 66 subjects classified according to burn severity (34 Severe and 32 LessSevere), with expression measurements organised across six time points. Missing visits were handled using feature–time mean imputation, with means estimated from training subjects within each validation partition.

Five-by-five subject-level nested cross-validation yielded 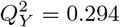, an AUROC of 0.827 and a balanced accuracy of 0.756 from pooled outer held-out predictions. Sensitivity and specificity were 0.794 and 0.719, respectively. Across the five outer folds, the selected number of PLS components ranged from 1 to 4 and the number of retained probes ranged from 2,047 to 2,212. Predictive results for both applications are summarised in Table 1.

**Table 1.** Predictive performance calculated from pooled outer held-out predictions in subject-level nested cross-validation. GSE68310 used training-fitted Tucker3 reconstruction with fixed ranks (4, 4, 3), whereas GSE182616 used training-derived feature– time mean imputation.

| Dataset | Q<sub>Y</sub><sup>2</sup> | AUROC | BA | Sens. | Spec. |
|---|---|---|---|---|---|
| GSE68310 | 0.670 | 0.971 | 0.931 | 0.945 | 0.917 |
| GSE182616 | 0.294 | 0.827 | 0.756 | 0.794 | 0.719 |

For full-cohort interpretation, component development selected *H* = 2. VIP-based selection retained 2,175 of the 50,599 probes, and the reduced model was refitted with the same two components. The selected-feature score representation is shown in Fig. 3A. Mean temporal *M*_2_ importance was highest at 4 hours, with a similar value at 2 hours, and lowest at 24 hours (Fig. 3B). Both panels summarize the full-cohort fitted models and are not held-out predictions.

**Figure 3.**
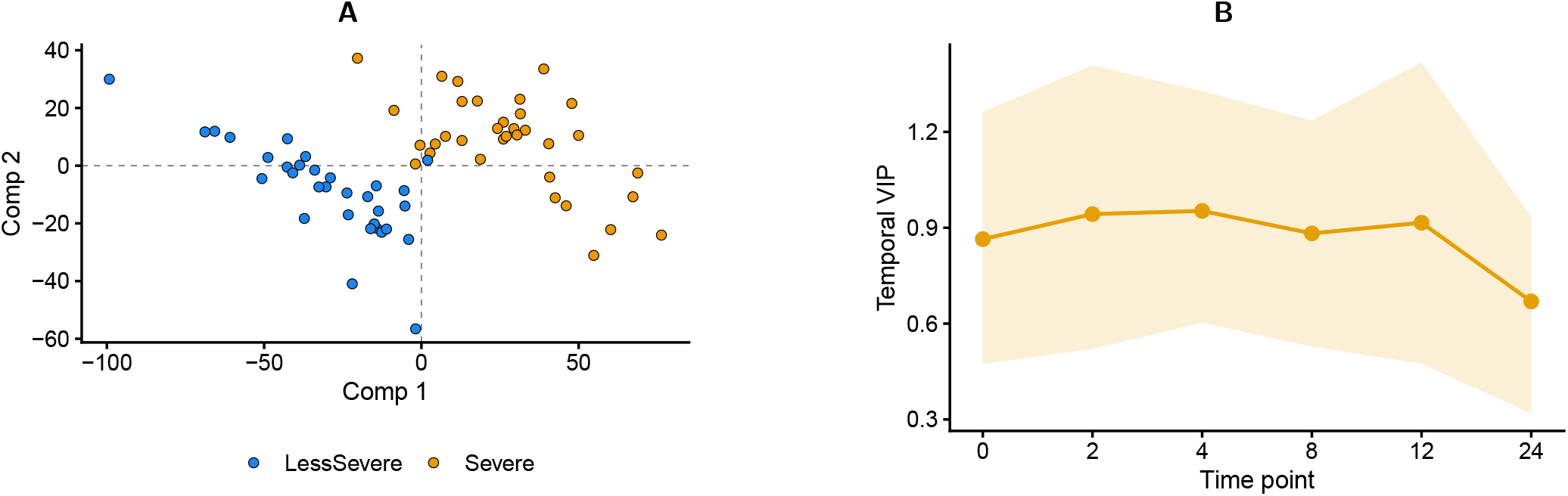
Full-cohort descriptive analysis of GSE182616 using feature–time mean imputation. (A) Sample scores after VIP-based feature selection and refitting on 2,175 probes. (B) Mean temporal *M*_2_ profile across selected probes; shading denotes the interquartile range across probes. Both panels describe fitted full-cohort models rather than held-out predictive performance. **Alt text:** Panel A plots full-cohort PLS-DA scores by burn-severity class. Panel B plots mean temporal *M*_2_ importance with interquartile shading, highest at four hours and lowest at 24 hours.

Functional enrichment identified no significant GO Biological Process terms, two Reactome pathways and one Hallmark gene set. The enriched Hallmark TNF*α* signaling via NF-*κ*B gene set included 42 selected genes (adjusted *P* = 1.88 × 10^−4^).

### Additional evaluations

We compared four VIP-based selection strategies (Supplementary Section S12 and Table S6). *M*_1_ retained the fewest probes and showed the highest fold-to-fold Jaccard similarity in both datasets (0.665 in GSE68310 versus 0.584 for *V* ∩ *M*_2_). In GSE68310, it also achieved the highest observed 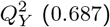, AUROC (0.974) and balanced accuracy (0.959), selecting a median of 500 probes per fold versus 1,122 for *V* ∩ *M*_2_. In GSE182616, *V* ∩ *M*_2_ yielded the highest 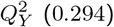 and AUROC (0.827), whereas *M*_1_ showed the highest balanced accuracy (0.758) and selection stability (Jaccard 0.323).

In the GSE68310 reconstruction benchmark, mean imputation had lower pooled RMSE at every masking level, although Tucker3 had lower RMSE in 55.47% of individual subject–mask comparisons. In simulations, Tucker3 improved reconstruction when shared longitudinal structure was informative, particularly at 10–20% masking, but could become unstable when few visits remained observed (Supplementary Sections S10–S11 and Figs. S1–S2).

## Discussion

The two applications illustrate TensorPLS on longitudinal transcriptomic datasets with different cohort sizes, sampling schedules and missing-visit patterns. The reconstruction benchmarks caution against assuming that Tucker3 is always preferable to feature–time mean imputation. Mean imputation had lower pooled RMSE in GSE68310, whereas Tucker3 improved reconstruction for synthetic data with informative shared longitudinal structure. At higher missingness, accuracy could deteriorate when too few visits remained to estimate reliable subject-specific coordinates. Imputation should therefore be chosen according to data structure and reconstruction performance rather than model complexity alone.

VIP-based selection revealed trade-offs between predictive performance, feature-set size and stability. *M*_1_ selected smaller, more stable probe sets in both datasets and gave the highest observed predictive metrics in GSE68310. However, *V* ∩ *M*_2_ achieved the highest 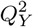 and AUROC in GSE182616. These differences were descriptive, do not establish a generally superior strategy and require evaluation on additional independent cohorts.

Nested cross-validation assessed held-out predictive performance, whereas full-cohort score plots and VIP profiles were descriptive. Enrichment provided biological context for selected genes, while temporal *M*_2_ profiles showed when their associated probes were important for discrimination. In GSE68310, interferon- and cell-cycle-associated probes peaked at different sampling times. Such importance does not indicate pathway activation or the direction of gene-expression changes.

This distinction between predictive evaluation and temporal interpretation also helps position TensorPLS relative to other approaches for longitudinal omics analysis. While tensorOmics fits tensor-native models and supports multi-block integration (Kodikara et al., 2026), TensorPLS applies matrix-based PLS-DA after subject-mode unfolding. Its emphasis is on integrating the analytical steps within a single workflow: during nested cross-validation, imputation, component tuning and VIP-based feature selection are performed within training partitions to assess predictive performance, while separate full-cohort models provide structured VIP profiles linked to functional enrichment for temporal interpretation. The two approaches are therefore complementary.

Both applications used processed GEO expression matrices, so the reported predictive performance is conditional on these deposited data. In GSE68310, enrolment year and recorded antiviral treatment differed between classes and may have contributed to discrimination. However, discrimination was retained in subgroup and cross-season analyses (Supplementary Sections S8–S9), although residual confounding cannot be excluded. Clinical characteristics and visit availability may also affect GSE182616. The two datasets address different clinical questions, but both show what TensorPLS brings to the analysis: a way to assess predictive performance and identify molecular features worth a closer look at different time points.

## Data availability

The processed gene-expression data and associated metadata analysed in this study are publicly available through the NCBI Gene Expression Omnibus under accession numbers GSE68310 and GSE182616.

For GSE68310, subject-level respiratory viral diagnoses were obtained from the supplementary phenotype data accompanying the GEO accession.

## Code availability

The TensorPLS R package is freely available in the TensorPLS GitHub repository, with a Zenodo record at doi:10.5281/zenodo.23674924.

Reproducibility code is available in the tensorPLSPaper GitHub repository.

## Author contributions

AG developed the TensorPLS R package, designed and implemented methodological extensions to the original framework, designed and performed the computational analyses, prepared the visualisations, and drafted the manuscript.

AC and LBN conceived the original research idea, developed the foundational methodology and initial computational scripts that served as the basis for TensorPLS, and contributed to the conceptualisation and methodological design of the study. They also supervised the work and reviewed and edited the manuscript. All authors reviewed and approved the final manuscript.

## Acknowledgments

We acknowledge the use of the Garnatxa high-performance computing cluster at I2SysBio (University of Valencia–CSIC).

## References

L. Balzano-Nogueira, R. Ramirez, T. Zamkovaya, J. Dailey, A. N. Ardissone, S. Chamala, J. Serrano-Quílez, T. Rubio, M. J. Haller, P. Concannon, M. A. Atkinson, D. A. Schatz, E. W. Triplett, and A. Conesa. Integrative analyses of TEDDY omics data reveal lipid metabolism abnormalities, increased intracellular ROS and heightened inflammation prior to autoimmunity for type 1 diabetes. Genome Biology, 22(1): 39, 2021. doi: 10.1186/s13059-021-02262-w.

M. Barker and W. Rayens. Partial least squares for discrimination. Journal of Chemometrics, 17(3):166–173, 2003. doi: 10.1002/cem.785.

R. Bro. Multiway calibration. multilinear PLS. Journal of Chemometrics, 10(1):47–61, 1996. doi: 10.1002/(SICI)1099-128X(199601)10:1⟨47::AID-CEM400⟩3.0.CO;2-C.

A. Conesa, J. M. Prats-Montalbán, S. Tarazona, M. J. Nueda, and A. Ferrer. A multiway approach to data integration in systems biology based on Tucker3 and N-PLS. Chemometrics and Intelligent Laboratory Systems, 104(1):101–111, 2010. doi: 10.1016/j.chemolab.2010.06.004.

S. Favilla, C. Durante, M. Li Vigni, and M. Cocchi. Assessing feature relevance in NPLS models by VIP. Chemometrics and Intelligent Laboratory Systems, 129:76–86, 2013. doi: 10.1016/j.chemolab.2013.05.013.

J. W. Keyloun, R. Campbell, B. C. Carney, R. Yang, S.-A. Miller, L. Detwiler, A. Gautam, L. T. Moffatt, R. Hammamieh, M. Jett, J. W. Shupp, and SYSCOT study group. Early transcriptomic response to burn injury: Severe burns are associated with immune pathway shutdown. Journal of Burn Care & Research, 43(2):306–314, 2022. doi: 10.1093/jbcr/irab217.

S. Kodikara, B. Lu, S. Wang, and K.-A. Lê Cao. tensorOmics: Data integration for longitudinal omics data using tensor factorisation. bioRxiv, 2026. doi: 10.64898/2026.02.10.705179.

T. G. Kolda and B. W. Bader. Tensor decompositions and applications. SIAM Review, 51(3):455–500, 2009. doi: 10.1137/07070111X.

A. Liberzon, C. Birger, H. Thorvaldsdóttir, M. Ghandi, J. P. Mesirov, and P. Tamayo. The molecular signatures database (MSigDB) hallmark gene set collection. Cell Systems, 1(6): 417–425, 2015. doi: 10.1016/j.cels.2015.12.004.

A. R. Taheriyoun, A. Ross, A. Safikhani, D. Soudbakhsh, and A. Rahnavard. Longitudinal omics data analysis: approaches and applications. Computational and Structural Biotechnology Journal, 31:301–315, 2026. doi: 10.1016/j.csbj.2026.01.001.

S. Varma and R. Simon. Bias in error estimation when using cross-validation for model selection. BMC Bioinformatics, 7: 91, 2006. doi: 10.1186/1471-2105-7-91.

S. Wold, M. Sjöström, and L. Eriksson. PLS-regression: a basic tool of chemometrics. Chemometrics and Intelligent Laboratory Systems, 58(2):109–130, 2001. doi: 10.1016/S0169-7439(01)00155-1.

G. Yu, L.-G. Wang, Y. Han, and Q.-Y. He. clusterProfiler: an R package for comparing biological themes among gene clusters. OMICS: A Journal of Integrative Biology, 16(5): 284–287, 2012. doi: 10.1089/omi.2011.0118.

Y. Zhai, L. M. Franco, R. L. Atmar, J. M. Quarles, N. Arden, K. L. Bucasas, J. M. Wells, et al. Host transcriptional response to influenza and other acute respiratory viral infections –a prospective cohort study. PLOS Pathogens, 11(6):e1004869, 2015. doi: 10.1371/journal.ppat.1004869. URL https://doi.org/10.1371/journal.ppat.1004869.

