## Supplementary Information for "TensorPLS: an R package for exploring molecular and temporal patterns in longitudinal omics data using PLS-DA"

#### Supplementary Methods

##### S1 GSE68310 data and model settings

The sensitivity analyses focused on GSE68310, comprising 109 subjects (73 influenza-positive and 36 positive for other respiratory viruses). Influenza-positive subjects included coinfections. The expression tensor contained 47,162 probes at Day0, Day2, Day4 and Day6, with 21 missing subject–time profiles (990,402 missing entries). Subject-level PCR and enrolment metadata were matched to the tensor identifiers.

The primary analysis used five outer and five inner cross-validation folds. Component counts $H = 1, \ldots, 10$ were evaluated using inner $Q^2_Y$, with a fixed `VIP2D_intersect_M2` feature-selection strategy and 99th-percentile threshold. Tucker3 imputation used ranks $(4, 4, 3)$, ridge coefficient 0.001, convergence tolerance $10^{-6}$, at most 2,000 iterations, HOSVD initialization and projection tolerance $10^{-10}$. The random seed was 123. The same Tucker3 settings, component candidates and VIP selection criteria were used for cross-season validation.

##### S2 Subgroup performance assessment

Performance was assessed separately for enrolment year (2009 or 2010) and infection complexity (monoinfection or coinfection, defined as more than one PCR-detected virus). For each subgroup, AUROC was recalculated using the continuous outer held-out predictions from the primary nested cross-validation. No models were refitted.

Sensitivity and specificity were calculated using the fixed threshold of 0.5; balanced accuracy (BA) was their mean. In both subgroup analyses, AUROC 95% percentile intervals were obtained from 2,000 outcome-stratified bootstrap resamples (seed 123), keeping predictions fixed. These intervals do not capture model-fitting or fold-selection variability.

##### S3 Cross-season validation

Two models were evaluated: one trained on 2009 and tested on 2010, and the other trained on 2010 and tested on 2009. Within each training year, five outcome-stratified inner folds were generated with `make_tensorpls_folds()` (seed 123). Tucker3 imputation, scaling, component tuning, VIP-based selection and model refitting were carried out using training subjects only. The resulting model was then applied to the held-out year.

AUROC, sensitivity, specificity and BA were calculated as described above. For each test year, $Q^2_Y$ used the training-year mean outcome as its reference:

$$Q^2_Y = 1 - \frac{\sum_{i \in \text{test}} (y_i - \hat{y}_i)^2}{\sum_{i \in \text{test}} (y_i - \bar{y}_{\text{train}})^2}.$$

#### S4 Real-data missing-visit reconstruction

We compared Tucker3 reconstruction and feature–time mean imputation using GSE68310. The experiment used the same five subject-level train/test partitions as the primary analysis. In each partition, both imputation methods were fitted on the training subjects and then evaluated on the held-out subjects.

To measure reconstruction accuracy, we artificially removed entire visits that were originally available in the test data. We tested five masking levels (5%, 10%, 20%, 30% and 40%), each repeated with three random seeds (123, 124 and 125). At least one visit was kept for each subject. Visits already missing in the original dataset were excluded from the accuracy calculation because their true measurements were unavailable.

Both methods reconstructed the same masked visits using parameters estimated from the training subjects. Mean imputation used the training feature–time means, whereas Tucker3 additionally used the remaining observed visits of each test subject to estimate subject-specific latent coordinates.

The reconstructed values were compared with the original measurements of the artificially removed visits. For each masking level and method, reconstruction accuracy was summarised using the pooled root mean squared error (RMSE) across the five folds and three repetitions:

$$\mathrm{RMSE}_{\mathrm{pooled}} = \sqrt{\frac{\sum_c \mathrm{SSE}_c}{\sum_c n_c}},$$

where $\mathrm{SSE}_c$ is the sum of squared reconstruction errors and $n_c$ is the number of evaluated entries in masking case $c$.

We also calculated RMSE separately for each subject with at least one artificially masked visit, recording how often either method achieved lower error and the number of visits remaining observed. These subject–mask evaluations are repeated observations rather than independent biological replicates.

For each masking level, the sums of squared errors (SSE) and evaluated entry counts were aggregated across masking repetitions for each subject. We then calculated 95% bootstrap percentile intervals for pooled RMSE using 2,000 resamples of subjects within each held-out fold (seed 123). The fitted models and masking patterns were kept fixed.

#### S5 Synthetic missing-visit reconstruction

We used simulated longitudinal data to examine how reconstruction accuracy varied with the underlying data structure, individual variation and measurement noise. Three scenarios were considered, each with three independently generated datasets containing 109 subjects, 1,000 features and four equally spaced time points.

For each feature, we first generated a mean temporal profile. Individual variation was then introduced through subject-specific offsets, linear trends and Gaussian-shaped temporal responses. Features were assigned to modules sharing the same underlying subject-level response parameters, with feature-specific coefficients controlling their contribution to each measurement.

Gaussian noise was added to the generated signal:

$$X_{ift} = L_{ift} + \varepsilon_{ift}, \qquad \varepsilon_{ift} \sim \mathcal{N}(0, \sigma^2),$$

where $L_{ift}$ is the noise-free signal and $X_{ift}$ is the simulated measurement for subject $i$, feature $f$ and time $t$.

The three scenarios were designed to represent different conditions. Scenario A contained strong individual variation shared across features within a single module. Scenario B retained this shared structure but introduced weaker individual variation and greater noise. Scenario C used 20 modules with more heterogeneous temporal responses. The main simulation parameters are summarised in Table S1.

Table S1: Main parameters used to generate the synthetic datasets. SD denotes standard deviation.

| Parameter | A | B | C |
|---|---|---|---|
| Modules | 1 | 1 | 20 |
| Individual offset SD | 0.4 | 0.032 | 0.4 |
| Response amplitude SD | 1.0 | 0.08 | 1.0 |
| Individual slope SD | 0.5 | 0.04 | 0.5 |
| Peak-position SD | 0.04 | 0.04 | 0.25 |
| Peak-width range | 0.35 | 0.35 | 0.12–0.35 |
| Noise SD | 0.3 | 0.6 | 0.3 |

Each dataset was divided into five subject-level train/test folds. Tucker3 and feature–time mean imputers were fitted using the complete training data. In the held-out subjects, entire visits were artificially masked at proportions of 10%, 20%, 30% and 40%, with three repetitions per level. At least one visit remained observed for each subject.

Both methods reconstructed the same masked visits without refitting on the test subjects. Tucker3 used the fixed parameters described in Section S1.

Reconstruction accuracy was evaluated against both the simulated noisy measurements and the underlying noise-free signal. Pooled RMSE was calculated as described in Section S4. Only cases with valid reconstructions from both methods were included in direct comparisons, while fitting failures were recorded separately. Subject-level RMSE was also calculated to examine the distribution of reconstruction errors.

#### S6 VIP strategy comparison and feature-selection stability

We compared predictive performance and feature-selection stability across four VIP-based strategies: the unfolded cumulative VIP representation ($V$), its time-averaged summary ($M_1$), its component-aggregated temporal summary ($M_2$), and the intersection $V \cap M_2$.

Each strategy was evaluated using subject-level nested cross-validation with five outer and five inner folds. Within each dataset, the same fold assignments, imputation method and candidate component counts ($H = 1, \ldots, 10$) were used across strategies. Each VIP strategy was evaluated separately using a fixed 99th-percentile threshold, while the number of PLS components was tuned independently within each outer-training partition.

For each candidate $H$, imputation, preprocessing, VIP-based feature selection and model refitting were performed using the inner-training subjects. The resulting predictions on the inner-validation subjects were pooled to calculate $Q_Y^2$, which was used to select $H$. The selected configuration was then refitted on the complete outer-training partition, including re-estimation of the selected feature set, and applied to the held-out outer-test subjects. This procedure produced one out-of-sample prediction per subject for each strategy.

Predictive $Q_Y^2$ was calculated by pooling squared prediction errors across all five outer folds:

$$Q_Y^2 = 1 - \frac{\sum_{k=1}^{5} \sum_{i \in \mathcal{T}_k} (y_i - \hat{y}_i)^2}{\sum_{k=1}^{5} \sum_{i \in \mathcal{T}_k} (y_i - \bar{y}_{-k})^2},$$

where $\mathcal{T}_k$ denotes the test subjects in outer fold $k$, $y_i$ is the observed binary response, $\hat{y}_i$ is its continuous held-out prediction, and $\bar{y}_{-k}$ is the mean response of the corresponding outer-training partition. Thus, $Q_Y^2$ was calculated from pooled prediction errors rather than by averaging fold-specific $Q_Y^2$ values. AUROC and balanced accuracy (BA) were calculated from the same pooled held-out predictions.

Feature-selection stability was assessed using the probe sets retained in the five outer-training partitions of each strategy-specific nested cross-validation result. For each pair of outer folds, we calculated the Jaccard index:

$$J(S_a, S_b) = \frac{|S_a \cap S_b|}{|S_a \cup S_b|},$$

where $S_a$ and $S_b$ denote the sets of molecular probes selected in folds $a$ and $b$. We report the mean Jaccard index across the ten distinct fold pairs and the median number of selected probes per fold. For the $V \cap M_2$ strategy used in the primary applications, we also calculated the union of probes selected in at least one fold and the intersection of probes selected in all five folds.

The primary $V \cap M_2$ cross-validation results were reused for this comparison; the other strategies were evaluated separately.

These comparisons describe predictive performance and feature-selection agreement under the same resampling partitions. They were not used to choose the selection strategy for the primary applications. Agreement across overlapping training partitions does not establish independent biological reproducibility, and no significance tests were performed on fold-pair Jaccard indices.

#### S7 Functional enrichment and temporal VIP analysis

**Probe-to-gene mapping and enrichment.** Selected microarray probes were mapped to Entrez gene identifiers using platform-specific annotations (GPL10558 for GSE68310 and GPL17077 for GSE182616). Only probes associated with a single valid Entrez identifier were retained for gene-level analysis. Identifiers were checked against org.Hs.eg.db, and duplicate gene identifiers were removed.

The gene background comprised all recognised Entrez identifiers represented among the analysed probes, rather than only the selected probes.

Over-representation analysis was performed using GO Biological Process, Reactome and MSigDB Hallmark gene sets (version 2026.1.Hs). Enrichment was assessed using the hypergeometric test, with Benjamini–Hochberg correction applied separately to each database. Gene sets containing 10–500 genes within the effective annotation background were considered, and adjusted $P < 0.05$ defined significance.

**Functional groups and temporal profiles.** For GSE68310, interferon-associated genes were defined as the union of the significant Hallmark interferon alpha and gamma response gene sets. Cell-cycle-associated genes were defined as the union of E2F targets and G2M checkpoint.

Genes shared between the two groups were excluded to obtain disjoint sets. The remaining selected probes were used as a reference group.

Temporal importance was evaluated using $M_2$ values from the original full-cohort all-feature PLS-DA model. For each functional group, the median and interquartile range of probe-level $M_2$ values were calculated at each sampling time.

### Supplementary Results

#### S8 Subgroup performance

AUROC ranged from 0.946 to 0.992 across the four subgroups (Table S2). Balanced accuracy was above 0.88 in all subgroups. Estimates were less precise for coinfections, which included only seven subjects in the Other class.

Table S2: Performance within GSE68310 subgroups based on the primary outer held-out predictions. Intervals are 95% stratified-bootstrap percentile intervals for AUROC.

| Subgroup | $N$ | Influenza | Other | AUROC [95% interval] | BA | Sens. | Spec. |
|---|---|---|---|---|---|---|---|
| 2009 | 56 | 31 | 25 | 0.992 [0.973–1.000] | 0.948 | 0.935 | 0.960 |
| 2010 | 53 | 42 | 11 | 0.946 [0.870–1.000] | 0.885 | 0.952 | 0.818 |
| Monoinfections | 78 | 49 | 29 | 0.984 [0.954–1.000] | 0.945 | 0.959 | 0.931 |
| Coinfections | 31 | 24 | 7 | 0.946 [0.851–1.000] | 0.887 | 0.917 | 0.857 |

#### S9 Cross-season performance

AUROC remained above 0.96 in both transfer directions (Table S3). Classification performance at the fixed threshold differed: when trained on 2010 and tested on 2009, the model achieved sensitivity of 1.000 but specificity of 0.640. The selected component count also differed between the two fits.

Table S3: Cross-season performance with Tucker3 imputation. The arrow indicates training to test year. AUROC intervals use 2,000 outcome-stratified bootstrap resamples of fixed test predictions.

| Training → test | Test $N$ | AUROC [95% interval] | BA | Sens. | Spec. | $Q^2_Y$ | $H$ | Probes |
|---|---|---|---|---|---|---|---|---|
| 2009 → 2010 | 53 | 0.963 [0.907–1.000] | 0.885 | 0.952 | 0.818 | 0.618 | 9 | 1,087 |
| 2010 → 2009 | 56 | 0.991 [0.970–1.000] | 0.820 | 1.000 | 0.640 | 0.637 | 6 | 1,299 |

#### S10 Real-data reconstruction performance

In GSE68310, feature–time mean imputation achieved lower pooled RMSE than Tucker3 at all five masking levels (Table S4; Fig. S1). All five Tucker3 training fits converged, and both methods were evaluated on the same 75 masking cases.

Although mean imputation performed better overall, Tucker3 achieved lower RMSE in 497 of 896 individual subject–mask evaluations (55.47%). The higher pooled error was largely associated with a small number of inaccurate reconstructions: the 5% of subject–mask evaluations with the largest Tucker3 squared errors accounted for 53.23% of the total Tucker3 squared error.

The proportion of evaluations favouring Tucker3 also varied with the number of remaining observed visits, increasing from 28.44% with one visit to 53.35% with two and 63.40% with three. These results suggest that the accuracy of subject-level Tucker3 reconstruction is sensitive to the information available for estimating individual latent coordinates. Because the same subjects

Table S4: Reconstruction accuracy on artificially masked visits in GSE68310. Pooled RMSE was calculated across five test folds and three masking repetitions per level. Lower values indicate better reconstruction accuracy.

| Masked visits | Mean RMSE | Tucker3 RMSE |
|---|---|---|
| 5% | 0.116 | 0.154 |
| 10% | 0.116 | 0.178 |
| 20% | 0.114 | 0.196 |
| 30% | 0.113 | 0.184 |
| 40% | 0.113 | 0.184 |

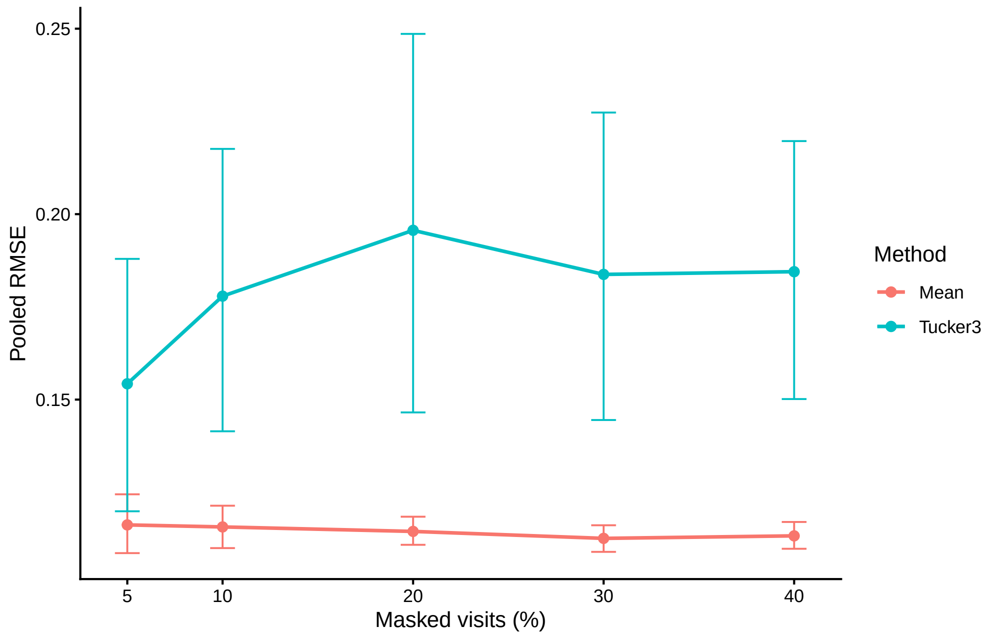


Figure S1: Reconstruction error across increasing levels of artificially masked visits in GSE68310. Lines show pooled RMSE for feature–time mean imputation and Tucker3 reconstruction. Error bars represent 95% percentile intervals from 2,000 within-fold subject-level bootstrap resamples. The fitted models and masking patterns were kept fixed. Lower RMSE indicates better reconstruction accuracy.

were evaluated under multiple masking conditions, these comparisons were treated as descriptive rather than independent observations.

#### S11 Synthetic reconstruction performance

Reconstruction accuracy differed across the three simulated scenarios (Table S5; Fig. S2).

In scenario A, Tucker3 achieved substantially lower RMSE than mean imputation when 10% or 20% of visits were masked. However, its pooled RMSE increased sharply at 30% and 40%, driven by large reconstruction errors in some held-out subjects.

In scenario B, both methods performed similarly at 10% masking, while mean imputation achieved lower RMSE at higher masking levels. In scenario C, Tucker3 showed a smaller but consistent improvement across all four levels. Similar patterns were observed when reconstruction errors were calculated against the underlying noise-free signal.

Table S5: Pooled RMSE for reconstruction of artificially masked visits in the three synthetic scenarios. Accuracy was evaluated against both noisy measurements and the underlying noise-free signal. Results include only masking cases with valid predictions from both methods. Lower RMSE indicates better reconstruction.

| | | | Noisy RMSE | | Latent RMSE | |
|---|---|---|---|---|---|---|
| Scenario | Masked | Pairs | Mean | Tucker3 | Mean | Tucker3 |
| A | 10% | 45 | 0.907 | 0.308 | 0.857 | 0.071 |
| A | 20% | 45 | 0.931 | 0.392 | 0.882 | 0.252 |
| A | 30% | 45 | 0.924 | 2.545 | 0.875 | 2.528 |
| A | 40% | 45 | 0.929 | 3.684 | 0.879 | 3.672 |
| B | 10% | 39 | 0.607 | 0.605 | 0.091 | 0.080 |
| B | 20% | 39 | 0.607 | 0.609 | 0.093 | 0.104 |
| B | 30% | 39 | 0.607 | 0.638 | 0.092 | 0.218 |
| B | 40% | 39 | 0.607 | 0.658 | 0.092 | 0.271 |
| C | 10% | 45 | 0.814 | 0.773 | 0.755 | 0.711 |
| C | 20% | 45 | 0.833 | 0.796 | 0.776 | 0.737 |
| C | 30% | 45 | 0.833 | 0.802 | 0.776 | 0.743 |
| C | 40% | 45 | 0.834 | 0.808 | 0.778 | 0.750 |

Noisy RMSE measures reconstruction error against simulated measurements containing noise, whereas latent RMSE measures error against the underlying noise-free signal.

Overall, 43 of the 45 Tucker3 training fits converged. The two non-convergent fits occurred in scenario B, leaving 516 valid paired masking cases out of 540 planned cases.

These results show that Tucker3 can improve reconstruction when the simulated data contain informative shared structure, but its accuracy may deteriorate when the remaining observations do not support stable subject-level reconstruction.

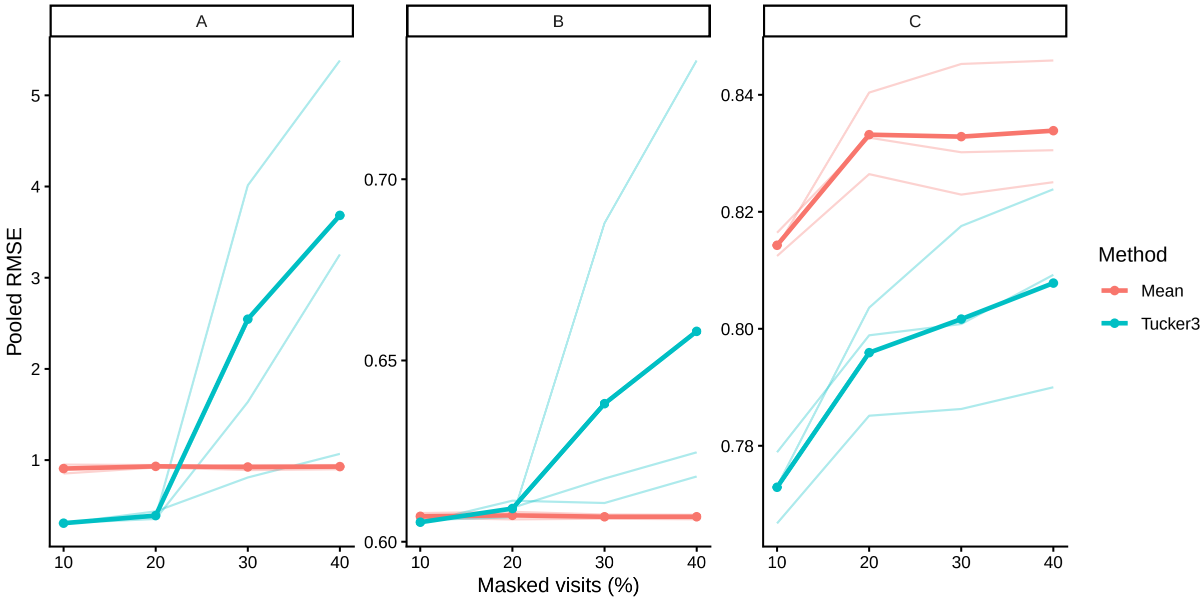


Figure S2: Reconstruction error across increasing levels of artificially masked visits in the three synthetic scenarios (A–C). Thick lines show pooled RMSE across valid paired evaluations, while thin lines show RMSE for each of the three independently simulated datasets. RMSE was calculated against the noisy measurements. Only masking cases with valid predictions from both methods were included. Lower RMSE indicates better reconstruction accuracy. Vertical axis scales differ across panels.

#### S12 Predictive performance and feature-selection stability

Predictive performance and feature-selection stability varied across VIP strategies and datasets (Table S6). $M_1$ produced the smallest median probe sets and the highest observed mean pairwise Jaccard in both datasets.

In GSE68310, $M_1$ achieved the highest $Q^2_Y$ (0.687), AUROC (0.974) and BA (0.959), while retaining a median of 500 probes per fold. Its mean Jaccard was 0.665, compared with 0.584 for $V \cap M_2$.

In GSE182616, $V \cap M_2$ achieved the highest $Q^2_Y$ (0.294) and AUROC (0.827). $M_1$ showed the highest observed BA (0.758) and mean Jaccard (0.323), with a median of 542 selected probes per fold.

Table S6: Predictive performance and probe-selection agreement across VIP strategies. $Q_Y^2$, AUROC and balanced accuracy (BA) were calculated from pooled outer held-out predictions. Probe counts are medians across the five outer folds, and Jaccard values are means over the ten distinct fold pairs. The all-feature model is omitted because it does not perform feature selection.

| Dataset | Strategy | Median probes | Mean Jaccard | $Q_Y^2$ | AUROC | BA |
|---|---|---|---|---|---|---|
| GSE68310 | $V$ | 1,485 | 0.565 | 0.664 | 0.970 | 0.931 |
| | $M_1$ | 500 | 0.665 | 0.687 | 0.974 | 0.959 |
| | $M_2$ | 1,304 | 0.567 | 0.664 | 0.969 | 0.917 |
| | $V \cap M_2$ | 1,122 | 0.584 | 0.670 | 0.971 | 0.931 |
| GSE182616 | $V$ | 2,684 | 0.248 | 0.285 | 0.824 | 0.756 |
| | $M_1$ | 542 | 0.323 | 0.262 | 0.826 | 0.758 |
| | $M_2$ | 2,538 | 0.259 | 0.250 | 0.803 | 0.728 |
| | $V \cap M_2$ | 2,194 | 0.244 | 0.294 | 0.827 | 0.756 |

For the $V \cap M_2$ strategy, 629 probes were selected in all five folds of GSE68310, among 1,847 probes selected in at least one fold. In GSE182616, 368 probes were selected in all five folds, among 6,084 selected at least once.

The $V \cap M_2$ strategy was used in the primary applications to illustrate how two VIP-based criteria can be combined to retain features satisfying both. It was not selected to maximise predictive performance or fold-to-fold stability. The observed differences therefore describe how predictive performance, feature-set size and selection agreement vary with the chosen VIP representation. These comparisons are descriptive and do not establish a statistically superior strategy.

#### S13 Functional enrichment and temporal VIP results

**Functional enrichment.** After probe-to-gene mapping and identifier validation, GSE68310 contained 921 selected genes among 23,306 background genes. GSE182616 contained 1,651 selected genes among 20,859 background genes.

The effective annotation backgrounds differed across databases. For GSE68310, 17,807 genes were represented in GO Biological Process and 11,168 in Reactome. For GSE182616, the corresponding backgrounds contained 16,877 and 11,089 genes.

In GSE68310, enrichment identified 306 significant GO Biological Process terms, 55 Reactome pathways and seven Hallmark gene sets. GSE182616 yielded no significant GO Biological Process terms, two Reactome pathways and one Hallmark gene set.

The significant Hallmark gene sets are reported in Table S7.

**Temporal importance of functional groups.** For GSE68310, the interferon and cell-cycle groups contained 83 and 81 selected probes, respectively, after excluding genes shared between the two groups. Two probes associated with shared genes were excluded, leaving 966 other selected probes as the reference group.

Using $M_2$ values from the all-feature model, interferon-associated probes reached their highest median importance at Day0 (3.08), whereas cell-cycle-associated probes peaked at Day4 (2.85). The remaining selected probes also showed their highest median importance at Day4 (2.62).

Table S7: Significant Hallmark gene sets identified by over-representation analysis. Overlap denotes the number of selected genes associated with each gene set. Adjusted $P$ values were calculated using the Benjamini–Hochberg procedure.

| Dataset | Hallmark gene set | Overlap | Adjusted $P$ |
|---|---|---|---|
| GSE68310 | Interferon alpha response | 48 | $1.14 \times 10^{-24}$ |
| | Interferon gamma response | 60 | $8.35 \times 10^{-18}$ |
| | E2F targets | 48 | $3.76 \times 10^{-10}$ |
| | G2M checkpoint | 41 | $1.08 \times 10^{-6}$ |
| | MYC targets V1 | 38 | $2.26 \times 10^{-5}$ |
| | MTORC1 signaling | 30 | 0.0166 |
| | Heme metabolism | 29 | 0.0261 |
| GSE182616 | TNF$\alpha$ signaling via NF-$\kappa$B | 42 | $1.88 \times 10^{-4}$ |

#### S14 Interpretation and limitations

The subgroup analyses showed that discrimination was not confined to one enrolment year or infection-complexity group. Cross-season validation provided a stricter assessment, with high AUROC in both transfer directions. However, specificity was lower when training on 2010 and testing on 2009, and clinical or technical factors may still have contributed to the observed discrimination.

The reconstruction experiments showed that Tucker3 was not consistently more accurate than feature–time mean imputation. Its advantages and limitations depended on the simulated data structure and on the information available to reconstruct held-out visits; the three synthetic scenarios were not designed to isolate individual generating parameters.

Finally, feature-selection agreement varied across datasets and VIP representations. The selected probe sets reflect the particular training partitions and selection criteria, and should be treated as candidates for further investigation rather than independently validated biomarkers.
